# Transcriptome analysis of *Medicago truncatula* Autoregulation of Nodulation mutants reveals that disruption of the SUNN pathway causes constitutive expression changes in a small group of genes, but the overall response to rhizobia resembles wild type, including induction of *TML1* and *TML2*

**DOI:** 10.1101/2023.01.19.524769

**Authors:** Elise L. Schnabel, Suchitra A. Chavan, Yueyao Gao, William L. Poehlman, F. Alex Feltus, Julia A. Frugoli

## Abstract

Nodule number regulation in legumes is controlled by a feedback loop that integrates nutrient and rhizobia symbiont status signals to regulate nodule development. Signals from the roots are perceived by shoot receptors, including a CLV1-like receptor-like kinase known as SUNN in the annual medic *Medicago truncatula*. In the absence of functional SUNN, the autoregulation feedback loop is disrupted, resulting in hypernodulation. To elucidate early autoregulation mechanisms disrupted in *SUNN* mutants, we searched for genes with altered expression in the loss-of-function *sunn-4* mutant and included the *rdn1-2* autoregulation mutant for comparison. We identified constitutively altered expression of small groups of genes in *sunn-4* roots, including higher levels of transcription factor *NF-YA2*, and in *sunn-4* shoots. All genes with verified roles in nodulation that were induced in wild type roots during the establishment of nodules were also induced in *sunn-4*, including, surprisingly, autoregulation genes *TML2* and *TML1*. Among all genes with a differential response to rhizobia in wild type roots, only an isoflavone-7-O-methyltransferase gene (Medtr7g014510) was found to be unresponsive in *sunn-4*. In shoot tissues of wild type, eight rhizobia-responsive genes were identified, including a MYB family transcription factor gene (Medtr3111880) which remained at a baseline level in *sunn-4*; three genes were found to be induced by rhizobia in shoots of *sunn-4* but not wild type. We also cataloged the temporal induction profiles of many small secreted peptide (MtSSP) genes in nodulating root tissues, encompassing members of twenty-four peptide families, including the CLE and IRON MAN families. The discovery that expression of *TML* genes in roots, a key factor in inhibiting nodulation in response to autoregulation signals, is also triggered in *sunn-4* in the section of roots analyzed suggests that the mechanism of TML regulation in *M. truncatula* may be more complex than published models.

## 1. Introduction

Legumes can grow on nitrate poor soil because of the ability to establish a symbiosis with nitrogen fixing soil bacteria. The legume rhizobia symbiosis, in which rhizobia inhabit the roots of legume plants and fix nitrogen from the atmosphere in exchange for carbon from photosynthesis, is an example of complex signaling between two very different species over both space (soil, root, shoot) and time. The time from rhizobial encounter in the soil to established nitrogen fixing nodules ranges from 8 to 20 days depending on species and conditions. Rhizobia secrete Nod factor in response to flavonoids exuded from legume roots; then, in the distal root area of emerged root hairs known as the maturation zone, the root hairs curl and entrap the bacteria, and calcium pulses triggered by the interaction rapidly begin altering plant gene expression. An infection thread develops around the dividing rhizobia and passes through the outer cortex to already dividing cells in the inner cortex, where the bacteria are released from infection threads into the developing nodule (reviewed in Ferguson et al., 2019; Oldroyd, 2013; Roy et al., 2020).

At each step in establishing symbiosis, there are checkpoints that must be cleared, as evidenced by the number of plant mutants in nodulation arrested at different developmental stages (Roy et al., 2020). Some of the “decisions” the plant must make include: Is this bacteria friend or foe? If friend, is this species a compatible friend and how is the bacteria allowed in without letting pathogens in as well? Are the correct sugars being made for passage through the thread? Is a particular thread headed to the primordia or should it be arrested? A large percentage of even compatible interactions are arrested in the outer cortex (Gage, 2004) for reasons not yet known. All of these “decisions” require molecular communication between the plant and microbe.

Once the plant has committed to the nodulation process, it controls the number of nodules that form and monitors the nitrogen output as there is an energy cost to the process of 12 to 16 g of carbon per gram of nitrogen fixed (Crawford et al., 2000), and intermittent or excess nitrogen offers little advantage to the plant. Multiple genes have been shown to control nodule number (see reviews Chaulagain and Frugoli, 2021; Roy et al., 2020) including genes in the autoregulation of nodulation (AON) pathway. This systemic pathway is initiated by the interaction of roots with rhizobia followed by transport of newly synthesized mobile peptide signals to a receptor complex in the shoot. In coordination with other pathways monitoring nutrient status, perception of the signal in the shoots causes changes in cytokinin and auxin flux and reduced transport of a mobile miRNA (*miR2111*) to roots which is proposed to allow accumulation of the nodulation inhibiting proteins that are the targets of the miRNA (reviewed in Chaulagain and Frugoli, 2021). The effects of AON can be detected in *M. truncatula* before nodules have fully developed, within 48 hours of inoculation (Kassaw and Frugoli, 2012), suggesting that AON signaling is happening simultaneously with nodule development.

In order to discover genes contributing to the developmental and nodulation phenotypes of AON mutants, we focused on early time points where AON and nodule initiation are happening simultaneously. We explored temporal gene expression differences between the sequenced wild type (A17) of the model indeterminate nodulating legume *Medicago truncatula* and mutants with lesions in two different genes in the AON pathway, the *SUPERNUMERARY NODULES* (*SUNN*) mutant *sunn-4* and the *ROOT DETERMINED NODULATION1* (*RDN1*) mutant *rdn1-2. SUNN* encodes a CLAVATA1-like leucine rich repeat receptor-like kinase, which is a key regulator of nodule number acting as a shoot receptor for mobile signaling CLE peptides induced in roots by rhizobia as well as mycorrhizae and nitrogen (Müller et al., 2019; Penmetsa et al., 2003; Schnabel et al., 2005). While all *sunn* alleles have short roots and auxin transport defects, the *sunn-4* allele has a stop codon very early in the coding sequence, and this null allele has a stronger hypernodulation phenotype than the original *sunn-1* allele, which harbors a kinase domain missense mutation (Schnabel et al., 2005; Schnabel et al., 2010). The *RDN1* gene encodes a hydroxyproline O-arabinosyltransferase that modifies one of the root-to-shoot signaling peptides, MtCLE12, enhancing transport and/or reception by the SUNN kinase (Kassaw et al., 2017). The *rdn1-2* allele has an insertion within the gene that greatly reduces the level of mature *RDN1* mRNA and affects the AON pathway upstream of SUNN (Kassaw et al., 2017; Schnabel et al., 2011).

In this work, we used a harvesting procedure described in (Poehlman et al., 2019) incorporating simultaneous inoculation of all plants in an aeroponic system and harvest of the zone of developing nodules tracked via root growth (summarized in Supplemental Fig 1). In our laboratory experience, nodulation progresses more uniformly and more rapidly in this system than on plates or in pots, and we presume this is due to the apparatus simultaneously spraying the aerosolized solution of rhizobia on all plants.

Using transcriptome data generated from the specific area of the root responding to our inoculation from these three genotypes, we identified a small collection of genes from those differentially expressed in both wild type and AON mutants responding to rhizobia which showed constitutively altered expression in root and shoot segments of the AON mutants. In examining the gene expression in nodulating root tissues during a time course covering the first 72 hours of interaction with rhizobia, as well as the shoots of these plants, we found that the AON mutants have a transcriptional response to rhizobia similar to wild type, including upregulation of many known nodulation pathway genes in roots and a small number of genes in the shoots. We found only two genes identified as responding to rhizobia in wild type that failed to also respond in *sunn-4*, one in roots and one in shoots, and these two genes displayed a moderated response in *rdn1-2*.

Unexpectedly, genes upregulated in both wild type and AON mutant roots included *TML1* and *TML2*, which encode nodulation inhibiting proteins in the AON pathway whose expression have been proposed to be increased by a SUNN-dependent decrease in *miR2111* levels in roots following nodule initiation using a split root system (Gautrat et al., 2019). Our observed induction of *TML1* and *TML2* in *sunn-4* root segments is not in agreement with a model in which increased *TML* levels in this area of the root at 48 hours post inoculation is sufficient to control nodule number, suggesting that additional factors dependent on SUNN function may be required for the later steps of AON or in other parts of the root.

## 2. Materials and methods

### 2.1. Tissue capture

Seeds of *Medicago truncatula* lines A17, *sunn-4*, and *rdn1-2* were scarified in concentrated sulfuric acid (93%–98%) by vortexing for 8 min in 50 mL sterile plastic conical tubes. The seeds were then rinsed in distilled water five times and imbibed for 3 to 4 h in water with gentle shaking at room temperature. Following imbibition, the seeds were plated by suspending over water on the lids of petri dishes and vernalized for 2 d in the dark at 4°C before being germinated overnight in darkness at 22°C. Seedlings were loaded onto an aeroponic growth apparatus described in (Penmetsa and Cook, 1997) and grown under a 14 h light/10 h dark cycle at 22°C in the described nodulation medium with no supplemental nitrogen. After 3 days of growth, samples of 20 plants per genotype (t = 0 h) were collected (3.5 h after initiation of daily light) and processed as described below and in Supplemental Fig 1A. For inoculations, 150 OD_600_ Units of *Sinorhizobium/Ensifer medicae* ABS7 resuspended in 40 to 100 ml of nodulation medium was added to the growth apparatus immediately after collection of the 0 h samples. Additional samples of 20 plants per genotype were collected 12, 24, 48, and 72 h later. Nodule primordia began the first cell divisions by 24 h after inoculation (Supplemental Fig 1B) and emerged nodules were visible on the root by 72 hpi.

Roots of ten plants per sample were measured to determine average root length for the following procedure. From the remaining ten plants, 2 cm segments representing the zone of development of the first nodules were collected. For 0 hpi samples, this region started 1 cm from the root tip, where the first full length root hairs were present. For later time points, the region of the first developing nodules was tracked by monitoring the progression of root growth away from the first interacting cells. The region was determined by calculating the average root growth since t=0 (average root length at t minus average root length at 0 h) and adding this distance to 1 cm, and 2 cm sections were collected upward from that position. This area is referred to as “root segments” throughout the text. Shoots, mid-hypocotyl and up, were collected from plants at the same time points. Collected tissue samples were placed into 1.5 ml tubes and stored at - 80°C.

### 2.2. RNA prep, libraries, and sequencing

RNA was purified from frozen samples by grinding in liquid nitrogen and using the RNAqueous Total RNA Isolation Kit (Invitrogen). Aliquots of RNA were analyzed for quality and concentration on an Agilent 2100 Bioanalyzer. RNA samples had RIN values between 8.3 and 9.9 for roots and 6.3 and 8.5 for shoots. Libraries for RNAseq were prepared and sequenced by Novogene Co., Ltd. (Beijing) from 100 to 1000 ng of total RNA using a stranded kit (Illumina TruSeq Stranded Total RNA Kit or NEB Next Ultra™ II Directional RNA Library Prep Kit for Illumina). The resulting data files contained paired end sequences (150 bp) ranged from 18,674,569 to 64,491,795 fragments.

### 2.3. Analysis of Gene Expression

The root data set consists of 75 libraries, including 60 libraries from this work (three replicates of five time points each for inoculated wild type A17, *sunn-4* and *rdn1-2* root segments, and uninoculated *sunn-4* root segments) and 15 libraries from uninoculated wild type A17 root segments generated in the same way and used in two manuscripts on new analysis algorithms (Gao et al., 2022; Poehlman et al., 2019). The shoot data set has 60 libraries (three replicates of five time points each for inoculated wild type A17, *sunn-4*, and *rdn1-2* and uninoculated A17), including 30 from this work for the two mutants and 30 from wild type previously reported (Gao et al., 2022). Read mapping and alignment data for each library is included in Supplemental Data Set 1. Among the genome v4.1 transcripts identified in our analyses were six that have been updated in v5, merging three pairs of transcripts (Medtr7g028557/Medtr7g028553, Medtr3g065345/Medtr3g065350, and Medtr0027s0200/Medtr0027s0180) into three genes (MtrunA17Chr7g0224711, MtrunA17Chr3g0110301, and MtrunA17Chr7g0240781, respectively). All the v4.1 transcripts are listed in the figures, but the v5 annotations were used for determining gene totals.

The data sets were processed with a DESeq2 pipeline using *Medicago truncatula* genome v4.1 as described in (Poehlman et al., 2019). We tested for differential expression at each time point using a cutoff of adjusted p-value <0.05 and minimum fold change of 2. Three pairwise comparisons of gene expression levels were performed at each timepoint (0, 12, 24, 48 and 72 hours post inoculation (hpi)) both for root segments and for shoots to create gene lists for further screening: (1) *sunn-4* (inoculated) versus A17 (inoculated), (2) *rdn1-2* (inoculated) versus A17 (inoculated), and (3) A17 (inoculated) versus A17 (uninoculated). Genes that were flagged as differential between the two 0 h datasets of A17 root segments, between the two 0 h datasets of A17 shoots, and between two 0 h datasets of *sunn-4* root segments were excluded from further analysis in those tissues to eliminate noise.

To identify genes with constitutively higher or lower expression in the AON mutants, genes from all three comparisons were further screened by assessing expression across all time points with heat map filtering and visual analysis of time course graphs. From 10,299 candidate genes in root segments and 749 in shoots, 32 and 49 genes were identified with consistently higher or lower expression in AON mutant roots and shoots, respectively.

Genes identified in root segments of A17 inoculated versus uninoculated (12, 24, 48, or 72 hpi) and identified as expressed at least two-fold lower in *sunn-4* inoculated versus A17 inoculated (12, 24, 48, or 72 hpi) were also further assessed with heat map filtering and visual analysis of time course graphs. Of 2155 candidate rhizobial response genes in A17 root segments, 477 were flagged by DeSeq2 with lower in expression in inoculated *sunn-4* root segments compared to A17; heat map analysis and visual analysis of time course graphs identified 3 of these genes with clearly reduced rhizobial response in *sunn-4* roots segments.

To identify rhizobial response genes in shoots, differentially expressed genes from all shoot comparisons were further screened with heat map filtering and visual analysis of time course graphs. From 749 candidate genes, 11 rhizobial response genes of shoots were identified.

Expression changes in selected genes were assayed by quantitative PCR using biological replicates independent of those used for RNAseq analysis. RNA was purified using the E.Z.N.A. Plant RNA Kit (Omega Bio-Tek) from sections of five roots. The iScript cDNA Synthesis Kit (Bio-Rad) was used to synthesize cDNA from 350 ng of RNA. Relative gene expression was assayed on the iQ5 system (Bio-Rad) using iTaq Universal SYBR Green Supermix (Bio-Rad). Expression levels (fold change) were determined by comparison to expression of control gene PI4K (Medtr3g091400).

Functional enrichment analysis was performed with the Medicago Classification Superviewer (http://bar.utoronto.ca/ntools/cgi-bin/ntools_classification_superviewer_medicago.cgi) using the default settings and a significance threshold of p<0.05. (Herrbach, et al. 2017).

### 2.4. Data access

The raw data underlying this manuscript is deposited at the National Center for Biotechnology Information under BioProjects PRJNA554677 (inoculated roots; control roots from mutants; inoculated and control shoots) and PRJNA524899 (control roots from wild type).

## 3. Results

### 3.1. Constitutively altered gene expression in *sunn-4* roots and shoots

We compared gene expression in *sunn-4* and *rdn1-2* to wild type (A17) to identify genes that were always different regardless of time or treatment in the AON mutants. Twenty-seven genes were found to have consistently higher (n=15) or lower expression (n=12) in *sunn-4* root segments compared to wild type (Fig. 1). Nine of these genes were similarly altered in *rdn1-2*, and an additional three were altered in *rdn1-2* only (including *rdn1* itself). Among the eight genes more highly expressed in both AON mutants compared to wild type root segments were *NF-YA2* (Medtr7g106450), a CAAT-binding transcription factor known to influence nodulation (Baudin et al., 2015; Laloum et al., 2014), and *MtSPL4* (Medtr2g014200), a SQUAMOSA Promoter Binding Protein-Like transcription factor. SPLs bind to SBP domain binding sites in promoters (Klein 1996). Reduced expression of a subset of *SPL*s in *Lotus japonicus* was observed when *miR156* was overexpressed in roots, and the authors hypothesize *miR156* directly or indirectly targets *ENOD40*, a gene important to nodule biogenesis (Wang 2015). The expression of a gene predicted to encode a 55 amino acid type II membrane protein of unknown function (Medtr2g090685) was lower in both AON mutants. The group of genes was determined to be enhanced for transcription factor (p = 0.012, Medicago Classification Superviewer) and transporter activities (p = 0.007).

**Fig 1.**
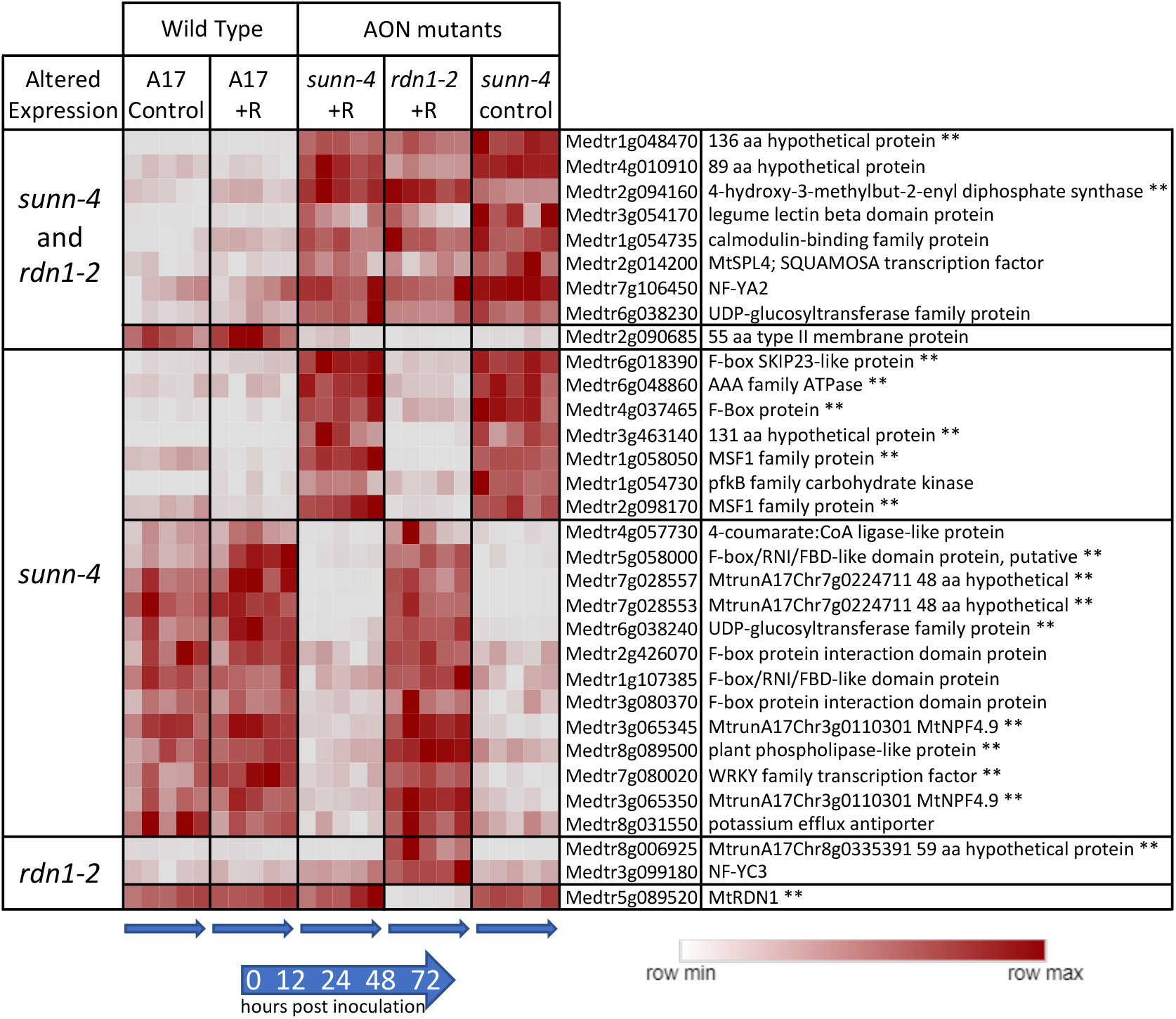
Genes with constitutively altered expression in roots of AON mutants *sunn-4* and *rdn1-2*. Heat map of average FPKMs of genes identified by DeSeq2 with altered expression levels in AON mutants compared to wild type (A17) that were consistent across all times and conditions (control = no rhizobia; +R = with rhizobia). Each row is independently scaled from minimum to maximum values; underlying data is in Supplemental Data Set 2. Expression of some genes was altered in both mutants, while for others the difference was only found in one mutant. Some genes had higher expression in the mutants and some had lower. The geneID (v4) and annotation is given. For five geneIDs, the annotation in v5 better matched the transcript structure; the v5 geneID is also given for these, with two pairs of geneIDs merged into two larger genes in v5. ** = also found to be similarly different in shoots of AON mutants (see Fig. 2).

Forty-one genes were found to have consistently higher (n=18) or lower expression (n=23) in *sunn-4* shoots compared to wild type (Fig. 2). Seventeen of these genes were similarly altered in *rdn1-2*, and an additional six were altered in *rdn1-2* only (including *rdn1* itself). Fourteen of the genes were also found among those higher (n=8) and lower (n=6) in roots of *sunn-4*.

**Fig 2.**
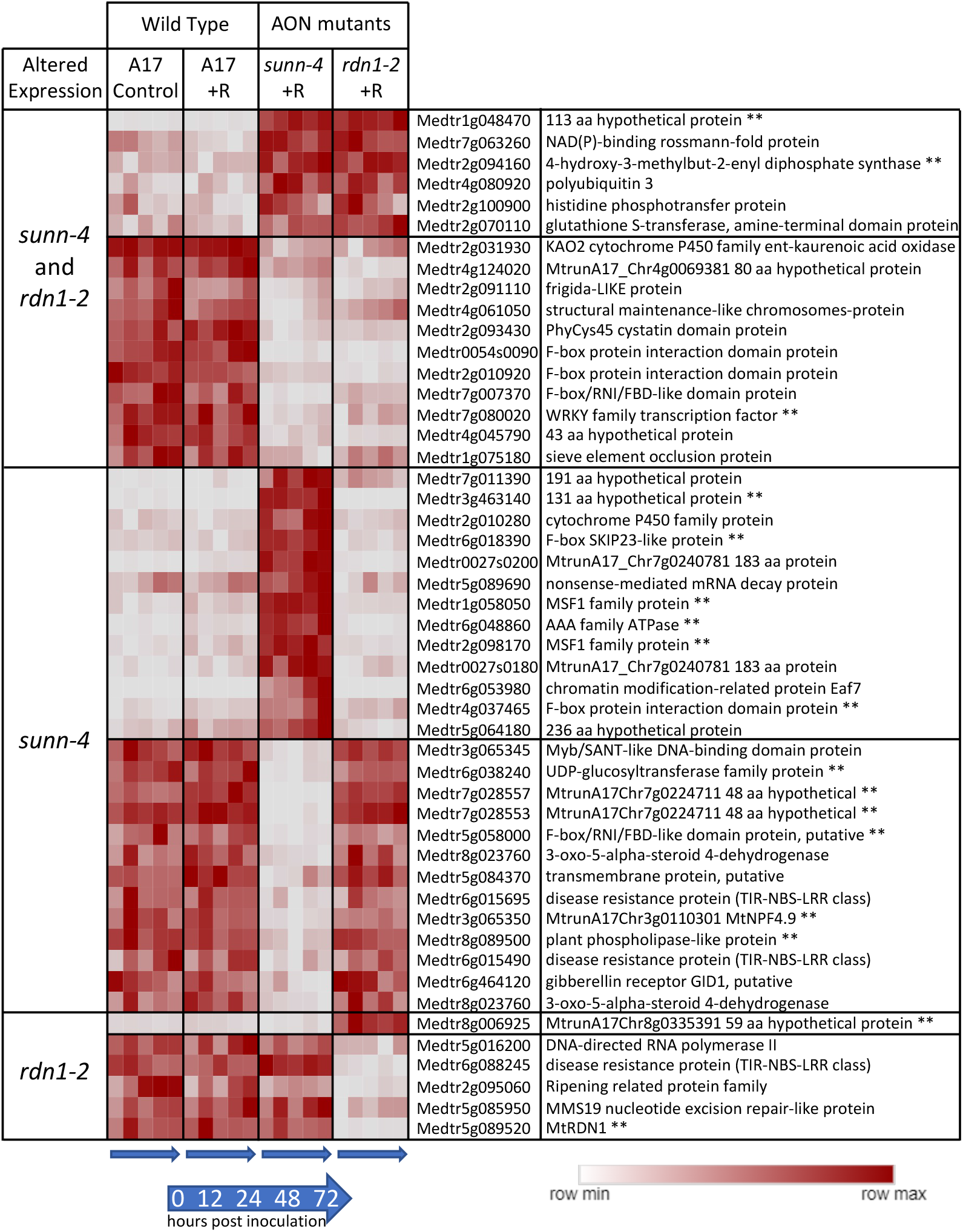
Genes with constitutively altered expression in shoots of AON mutants *sunn-4* and *rdn1-2*. Heat map of average FPKMs of genes identified by DESeq2 with altered expression levels in AON mutants compared to wild type (A17) that were consistent across all times and conditions (control = no rhizobia; +R = with rhizobia). Each row is independently scaled from minimum to maximum values; underlying data in in Supplemental Data Set 2. Expression of some genes was altered in both mutants, while for others the difference was only found in one mutant. Some genes had higher expression in the mutants and some had lower. The geneID (v4) and annotation is given. For seven geneIDs, the annotation in v5 better matched the transcript structure; the v5 geneID is also given for these, with two pairs of geneIDs merged into two larger genes in v5. ** = also found to be similarly different in roots of AON mutants (see Fig. 1).

### 3.2. Response of genes to rhizobia in roots in wild type and *sunn-4*

#### 3.2.1. Nodulation pathway genes

We assessed our dataset for the behavior of 207 functionally validated symbiotic nitrogen fixation genes ranging in roles from early nodulation signaling to nitrogen fixation (Roy et al., 2020). While not all these genes would be expected to respond to rhizobia, 56 of the 207 genes (27%) had increased expression after inoculation, while 2 showed lower expression in AON mutants (Fig. 3). *RDN1* showed consistently lower expression in *rdn1-2*, as had previously been shown for this mutant (Schnabel et al., 2011), and the synaptotagmin gene *MtSYT2* had decreased expression in AON mutants at the 72 hpi time point. *MtSYT2* encodes a synaptotagmin from a family of three in *M. truncatula* shown by localization and RNAi to be involved in formation of the symbiotic interface (Gavrin et al., 2017).

**Fig 3.**
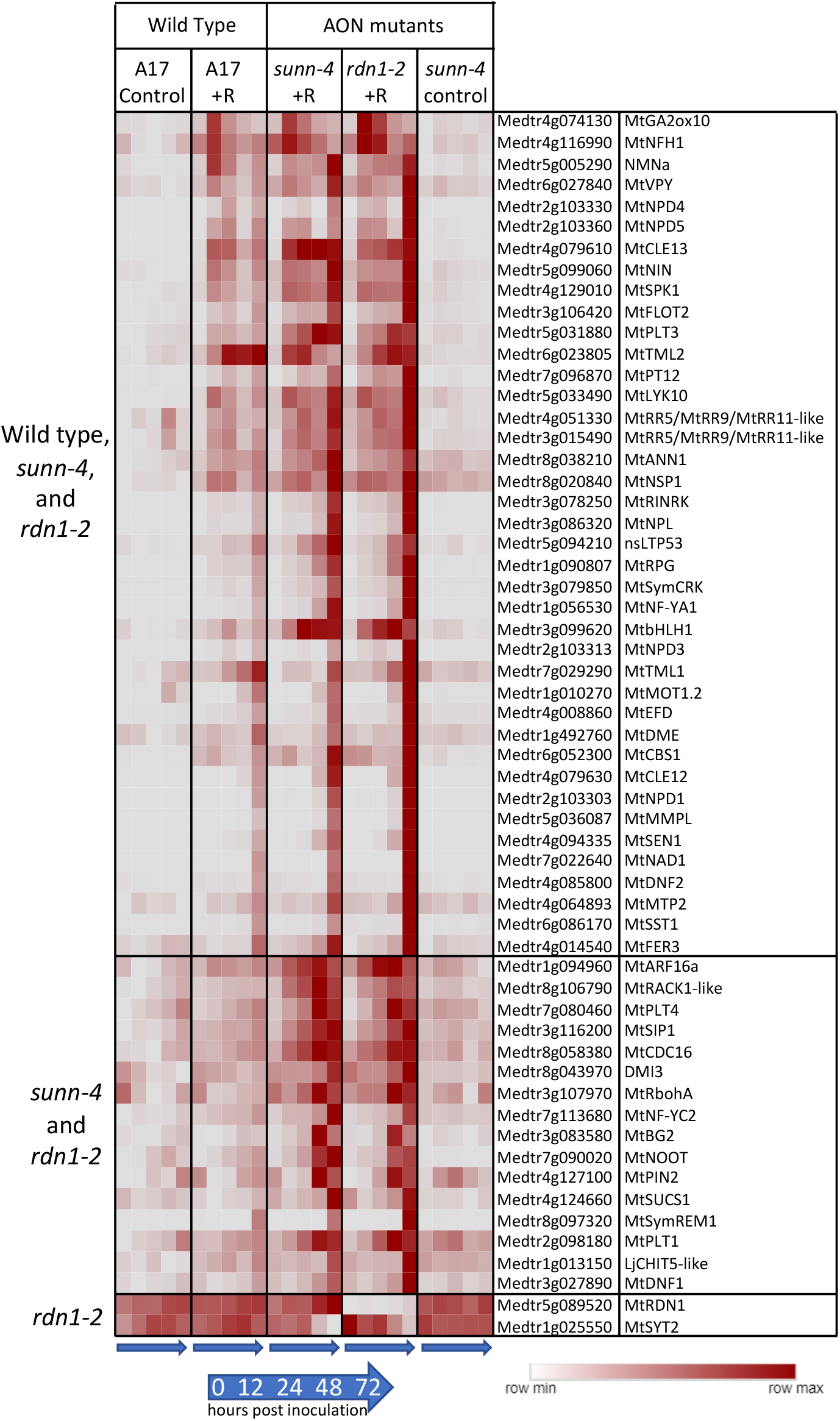
Rhizobia-induced expression of nodulation pathway genes in roots of wild type (A17) and/or AON mutants *sunn-4* and *rdn1-2*. Heat map of average FPKMs of 58 known nodulation genes with patterns of expression that changed with rhizobial inoculation (+R) or with genotype. Each row is independently scaled from minimum to maximum values; underlying data is in Supplemental Data Set 2. Induction was detected in all three lines for some genes (n=40) and for only the AON mutants for others (n=16). Two genes were altered in *rdn1-2* only.

Forty of the differentially expressed genes in Fig. 3 were induced by rhizobia in both wild type and AON mutants with most genes more highly induced in the mutants. Included among these genes induced in all genotypes are four Nodule PLAT domain proteins which are known to be expressed in nodules (Pislariu et al., 2019), but interestingly, unlike *MtNPD1* which showed increased expression only at 48 and 72 hpi, *MtNPD4* and *MtNPD5* had increased expression by 12 hpi and *MtNPD3* by 24 hpi. Induction of sixteen genes was only seen in the AON mutants and included five genes increasing by 12 hpi and eleven genes by 48 or 72 hpi. Included among genes induced early in AON mutants is the PLETHORA gene *MtPLT4*, which has been shown to be expressed in the central areas of nodule meristems (Franssen et al., 2015). *MtPLT1*, known to be expressed in peripheral areas of nodule meristems, was increased at 48 or 72 hpi timepoints.

#### 3.2.2. TMLs

Two genes, *TML1* and *TML2*, whose upregulation in roots is proposed to be a key part of the AON pathway downstream of SUNN, showed an unexpected response to rhizobia in AON mutants (Fig. 4). In wild type and in AON mutants, *TML1* and *TML2* RNA levels increased in response to rhizobia. *TML2* expression was induced by 12 hpi and peaked by 24 hpi in all three genotypes (Fig. 4A). For *TML1*, RNA levels started increasing around 24 hpi and continued to rise in wild type and *rdn1-2*; in *sunn-4, TML1* expression began to increase later (Fig. 4B). Given the rise in transcript abundance for both genes in *sunn-4*, we tailored the qPCR confirmation of the result to the times of induction. For *TML2* we chose to divide the interval before the increase into smaller fractions to verify our unexpected finding of increased RNA expression by qPCR, rather than repeat the entire time course. Since *TML1* expression rose later in the time course, we repeated the entire time course for *TML1*. The overall patterns of expression for *TML2* (Fig. 4C) and *TML1* (Fig. 4D) were similar to wild type in independent samples assayed, with wild type and *sunn-4* showing increased *TML2* by 8 hpi and *TML1* by 48 hpi.

**Fig. 4.**
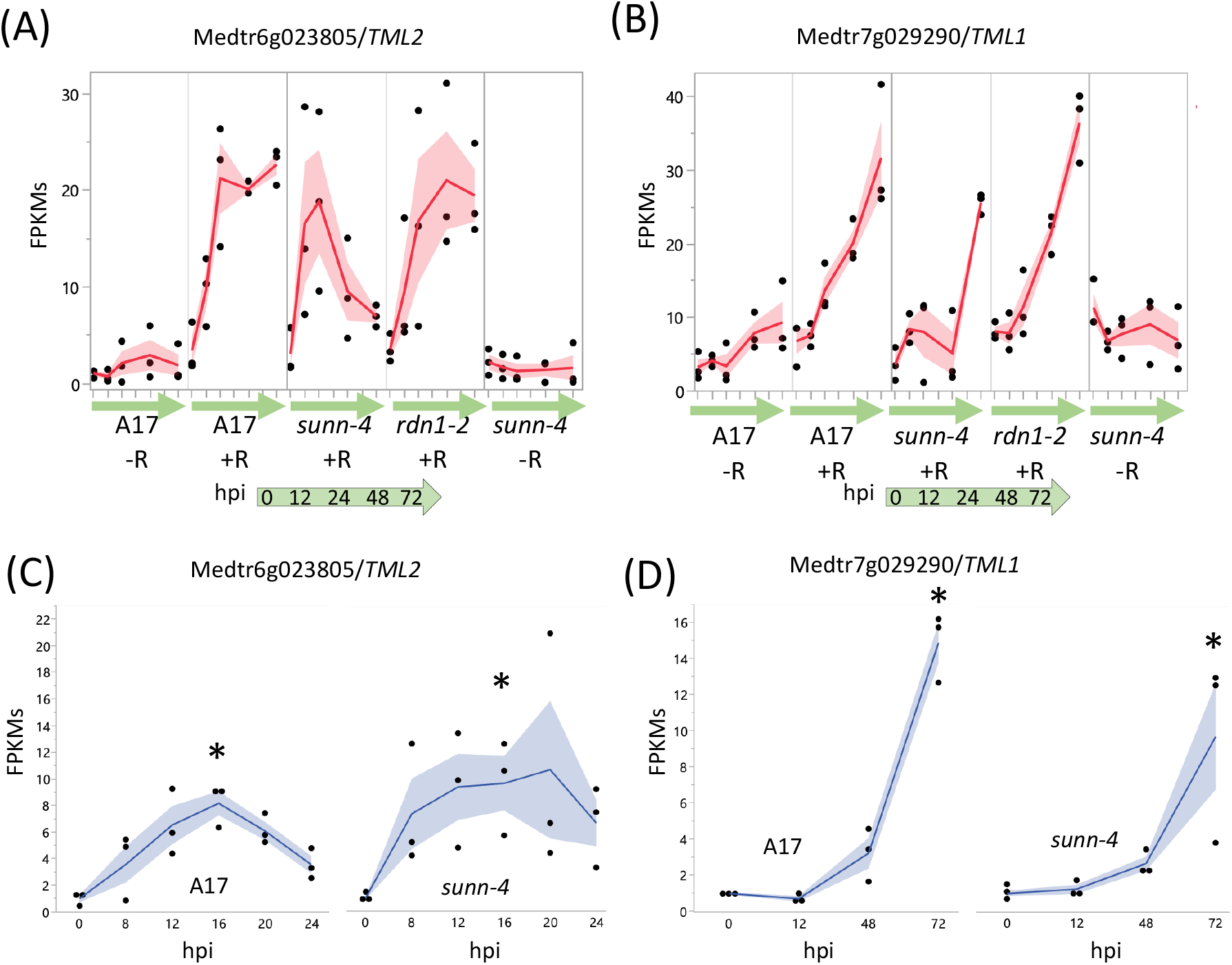
AON genes *TML2* and *TML1* are induced by rhizobia in both wild type and AON mutants. RNA-seq data shows early induction in response to rhizobia for *TML2* (A) and later induction for *TML1* (B). The FPKMs (black dots) and means (red lines) of three biological replicates are shown for time points 0 through 72 hours post inoculation (hpi) for uninoculated (-R; wild type A17 and *sunn-4*) and inoculated (+R; wild type, *sunn-4*, and *rdn1-2*) root segments. qPCR verified the induction of *TML2* (C) and *TML1* (D) in both A17 and *sunn-* 4. Expression levels were significantly higher in both wild-type and *sunn-4* at 16 hpi for *TML2* and at 72 hpi for *TML1* (*, p<0.05; Kruskal-Wallis test with Bonferroni correction). The relative expression of three biological replicates (black dots = data points; blue line = means) of these genes is shown. Shading shows the standard error of the mean.

#### 3.2.3. Genes unresponsive to rhizobia in *sunn-4* mutants

A screen for genes induced by rhizobia in wild type but not in *sunn-4* yielded three genes, one with increased expression over 72 hpi (Fig. 5) and two with increased expression by 12 hpi that was then reduced (Supplemental Fig. 2). Following tests of independent samples by qPCR, it was found that an isoflavone 7-O-methyltransferase gene (Medtr7g014510) showed consistently increasing expression in wild type over the 72 hours following inoculation but showed a much lower increase in *sunn-4*. For the other two genes (Medtr2g086390, a b-ZIP transcription factor and Medtr1g109600 a putative small signaling peptide), the induction that was observed in the RNAseq data was not found in the qPCR data (Supplemental Fig. 2).

**Fig. 5.**
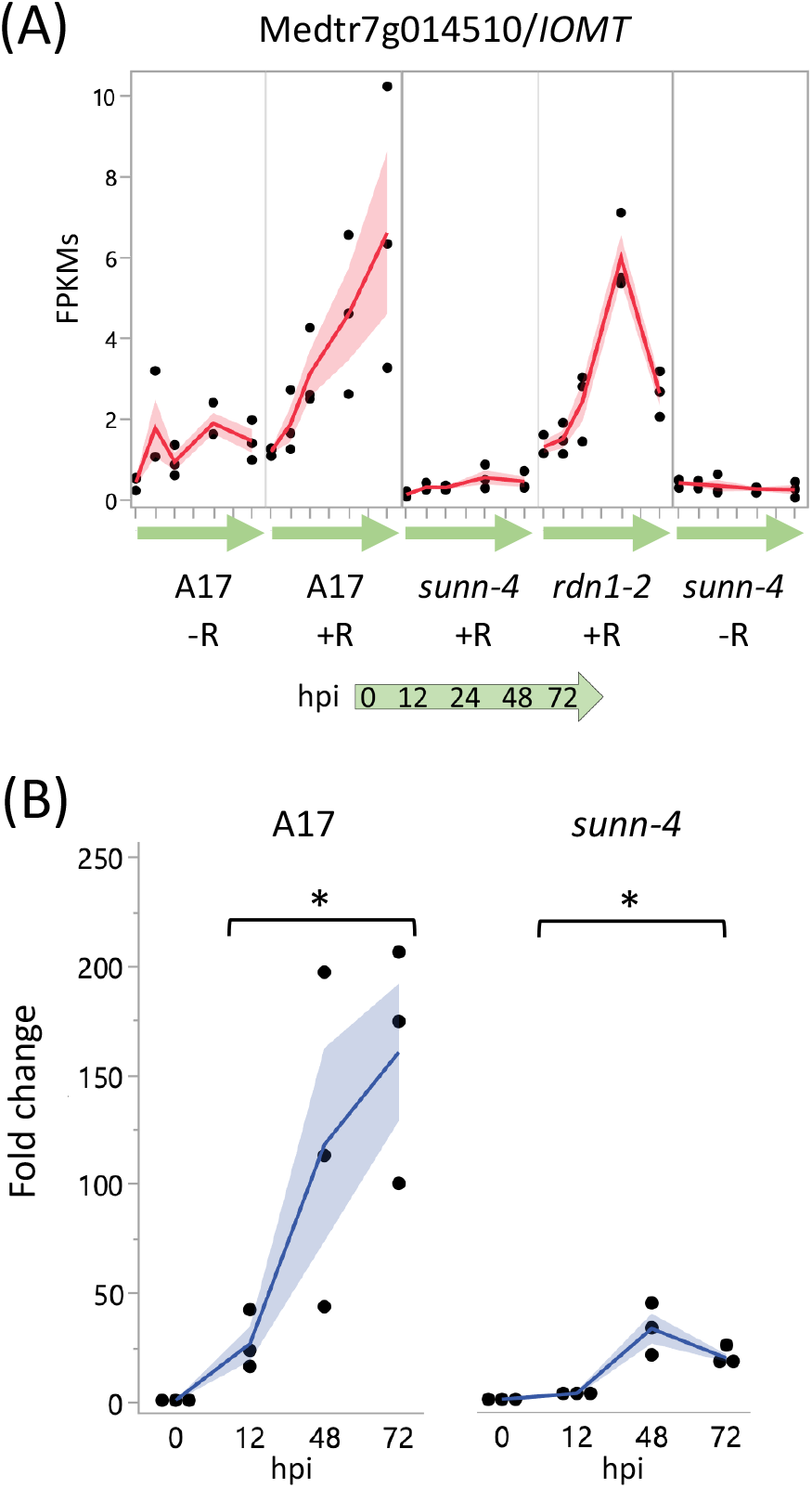
Strong induction of an Isoflavone 7-O-Methyltransferase (Medtr7g014510) in nodulating root segments of wild type is not seen in *sunn-4*. (A) FPKMs (black dots) and means (red lines) from three biological replicates for Medtr7g014510 from RNA-seq of wild type and the AON mutants *sunn-4* and *rdn1-2* over the first 72 hours post inoculation (hpi) with rhizobia (+R) compared to uninoculated controls (-R). (B) qPCR analysis of Medtr7g014510 in wild type and *sunn-4* showing expression levels relative to 0 h. Post inoculation times points were significantly higher than the 0 h samples, although the extent of induction was 3- to 9-fold less in *sunn-4* (*, p < 0.05; Kruskal-Wallis test). Shading is the standard error of the mean.

#### 3.2.4. Induction of Signaling Peptides genes in roots

Members of the CLE peptide gene family were previously identified as playing an important role in nodulation regulation (Mortier et al., 2010; Okamoto et al., 2009). We found 170 peptide-encoding genes from the *Medicago truncatula* Small Secreted Peptide Database (Boschiero et al., 2020) that showed a rhizobia-induced increase of expression in wild type and/or AON mutants (Table 1). Twenty-four peptide families are represented by these genes, including NCR peptides (n=46) and CLE peptides (n=10). Expression of some peptide genes began increasing by 12 hpi (n=23), such as seven plant defensins, while others were induced at later times (by 24 hpi, n=9; by 48 or 72 hpi, n=138), including all fourteen members of the IRON MAN peptide family (Grillet et al., 2018). Interestingly, although NCR peptides are known to accumulate in nodules to aid bacterial differentiation (Van de Velde et al., 2010), *NCR150* (Medtr6g466410) showed a transient increase in expression at 12 and 24 hpi (Supplemental Fig. 3), suggesting a role for NCR150 in a non-nodule tissue. High levels of *CLE12* and *CLE13* have been documented at 3 to 15 days post inoculation, with *CLE13* levels increasing earlier than CLE12 (Mortier et al., 2010); we found that *CLE13* is induced by 12 hpi, while *CLE12* levels do not begin to rise significantly until 48 to 72 hpi (Supplemental Fig. 4). Early expression of *CLE13* in AON mutants was as in wild type, although by 72 hpi levels in mutants were 2-fold higher, while induction of *CLE12* in AON mutants was stronger, consistent with the data in (Kassaw et al., 2017).

**Table 1.**
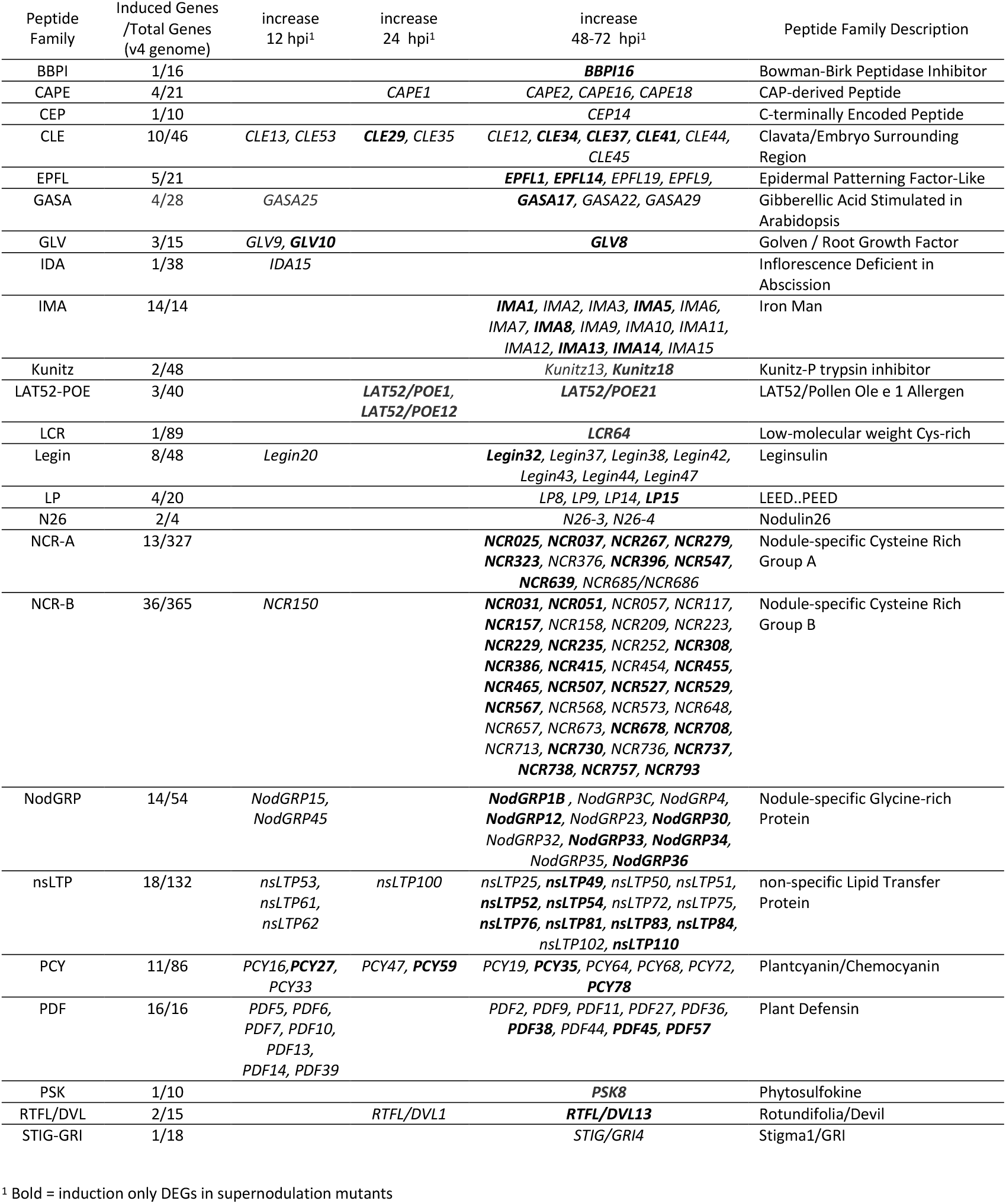
Small Secreted Peptide (MtSSP) genes induced by rhizobia in the maturation zone of wild type or autoregulation mutants *sunn-4* and *rdn1-2* during nodule development.

### 3.3. Rhizobial response in shoots

The systemic AON pathway is initiated by the interaction of roots with rhizobia followed by transport of newly synthesized mobile peptide signals to a receptor complex in the shoot. Perception of the signal in the shoots results in a signal sent to roots to inhibit further nodule development. From our DESeq2 pipeline, eleven genes were identified showing a rhizobial response in plant shoots (Fig. 6A). Seven genes were induced in both wild type and AON mutants over the first 72 hpi following rhizobia inoculation, while a single gene was induced in wild type but not *sunn-4* (Medtr3g111880, a predicted MYB family transcription factor), and three genes were induced in *sunn-4* but not in wild type (Medtr1g074990, Medtr2g435780, and Medtr4g121913). The seven genes increasing in all three lines include an IRON MAN Peptide gene (IMA11; Medtr4g026440), also found as one of fourteen IRON MAN genes induced by rhizobia in roots, and the gene for carotenoid cleavage dioxygenase 4a-6 (CCD4; Medtr5g025270).

**Fig 6.**
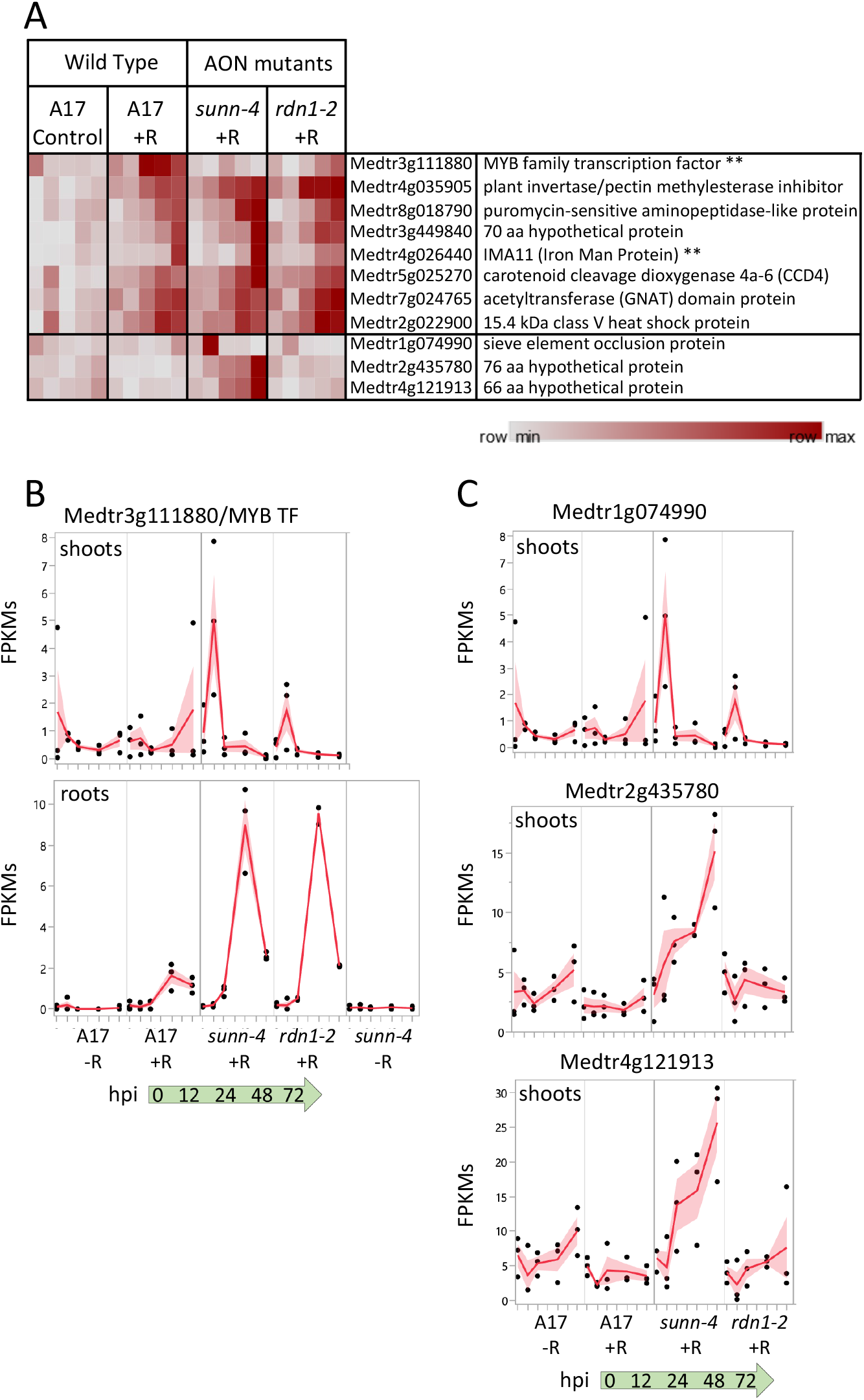
Gene expression induced by rhizobia in shoots. (A) Heat map of average FPKMs of genes showing increased expression during the first 72 hours of nodulation. Each row is independently scaled from minimum to maximum values; underlying data in in Supplemental Data Set 2. Eight genes were induced in wild type. Seven of these were also induced in AON mutants *sunn-4* and *rdn1-2*. Three additional genes were induced only in *sunn-4*. Two genes that were also induced in nodulating roots are indicated by “**”. Graphical representation of selected genes is shown in (B) and (C) with FPKMs (black dots) and their means (red lines) from three biological replicates. Shading is the standard error of the mean. (B) Transcription factor gene Medtr3g111880 was induced by 24 h in shoots of wild type but not *sunn-4*, while in roots expression was induced in all three lines. (C) Three genes induced in shoots of *sunn-4*.

The predicted MYB family transcription factor gene (Medtr3g111880) found to be uninduced in *sunn-4* increased expression in wild type by 24 hpi, but by 72 hpi there was still no increase detected in *sunn-4* (Fig. 6B). In the *rdn1-2* mutant, expression levels increased but on a slower time frame than in wild type, with the first increase seen at 48 hpi. This gene was also found to be induced by rhizobia by 48 hpi in roots, where the AON mutants both showed a stronger induction of expression than wild type.

A predicted sieve element occlusion protein gene (Medtr1g074990) showed an early transient pulse of expression in *sunn-4* that was weaker in *rdn1-2* and absent in wild type (Fig. 6C). Two genes encoding small proteins of unknown function (Medtr2g435780 and Medtr4g121913) showed increasing expression across the time course in *sunn-4* but no apparent increase in *rdn1-2* or wild type. None of these three genes showed a response in roots, with Medtr4g121913 expression restricted to shoots.

## 4. Discussion

### 4.1 Expression differences in AON mutants

Combining transcriptome data from wild type and two AON mutants, *sunn-4* and *rdn1-2*, we identified genes with altered expression in roots and shoots of the AON mutants. Using only the part of the root initially responding to rhizobia and following only that part of the root over time in synchronized plants was possible because of the aeroponic system we used to grow the plants. While the simultaneous aeroponic system has been used to generate a transcriptome in a previous study (Larrainzar et al., 2015) and the use of the area surrounding inoculation has been used on growth plates (Schiessl et al., 2019) the two methods have not been combined until this report.

Given the level of molecular communication between the plant and microbe and the critical role of SUNN in AON, we expected that disruption of *SUNN* would significantly impact transcriptomic responses to rhizobia. Intriguingly, most of the differences we found were constitutive and not specific to the rhizobial response, suggesting that the AON pathway is one of multiple signal transduction pathways affected by a mutation in SUNN. A microarray comparison of uninoculated *sunn-1* mutant against uninoculated *lss* mutants in which no *SUNN* transcript is produced (phenocopy of *sunn-4*) also showed that no *SUNN* expression resulted in less misregulation than mutant *SUNN* expression (Schnabel, et al 2010).

We identified 54 genes with constitutively altered expression in *sunn-4*. Among these are 14 genes with altered expression in both roots and shoots (8 higher, 6 lower), 13 in roots only (7 higher, 6 lower), and 27 in shoots only (10 higher, 17 lower). Expression of 23 of the genes was also altered in *rdn1-2*. An additional 7 genes had altered expression in *rdn1-2* only. Seven of these genes are predicted to encode proteins of unknown function (annotated as hypothetical proteins) with lengths ranging from 48 to 236 amino acids; six genes have only been identified in *M. truncatula* and the other gene has been described in clover. Also among the 54 genes are several members of much larger gene families, including two UDP glycosyltransferases (out of over 250 genes in the family), one legume lectin binding domain protein (out of 40) and one calmodulin-binding family protein (out of 8). The 4-hydroxy-3 methylbut-2-enyl diphosphate synthase (Medtr2g094160), which is more highly expressed in roots and shoots of both AON mutants, is an enzyme involved in the formation of isoprenoid-derived plant signaling compounds (Tarkowská and Strnad, 2018). Other genes misexpressed in *sunn-4* included some with suggested functionalities, such as F-box containing genes, transcription factors, and transporters. The collection of genes constitutively altered in *sunn-4* do not point to a known single pathway globally disrupted in these mutants but rather indicate multiple discreet differences.

A single gene demonstrated an attenuated rhizobial response in *sunn-4* roots. The isoflavone methyltransferase gene Medtr7g014510, encoding MtIOMT3 (Deavours et al., 2006), was upregulated in wild type and the *rdn1-2* mutant by 12 hours, whereas in *sunn-4* the upregulation was much weaker. This gene is induced in leaves infected with *Phoma medicaginis*, a known inducer of isoflavonoid synthesis (He and Dixon, 2000; Paiva et al., 1994). MtIOMT3 has been shown to be able to modify a variety of compounds including 6,7,4′-trihydroxyisoflavone, 7,3′,4′-trihydroxyisoflavone, genistein, glycitein, and dihydrodaidzein (He and Dixon, 2000). Isoflavones are endogenous regulators of auxin transport in soybean, and genistein production is also part of nodule development in soybean (Subramanian, et al. 2006). Since the *sunn-1* mutant has a defect in auxin transport (Van Noorden, G.E. et al. 2006), misregulation of MtIOMT3 is likely responsible for the defect.

### 4.2 Peptide Responses to Rhizobia

The breadth of the response observed in peptide-encoding genes (Table 1) reflects the ubiquitous nature of peptide function in plant roots (de Bang et al., 2017; Jeon et al., 2021; Kim et al., 2021), and members of multiple peptide-encoding gene families were identified in the nodulation response, including CLE peptides with a demonstrated role in nodulation regulation (Mortier et al., 2010; Okamoto et al., 2009). We found that the AON mutants showed a pattern of peptide-encoding gene response to rhizobia similar to wild type. The difference was an increased number of peptide-encoding genes induced, mostly at later time points, presumably due to the higher number of nodules present in the analyzed tissue. There were a few peptide expression patterns of note.

NCR peptides (Nodule-specific Cysteine-Rich Peptides), a large group of defensin-like antimicrobial peptides, are produced in nodules of *M. truncatula* and control rhizobial development (Guefrachi et al., 2014; Maróti, Downie, & Kondorosi, 2015). Interestingly, we found that whereas almost all NCR peptides were nodule-specific, a single NCR peptide gene (NCR150) was expressed at our earliest time point, before nodules had developed, and then was turned back off at nodule emergence. NCR150 has previously been shown to be one of the very few NCR peptides known to be expressed outside of nodules, as it was found in epidermal cells following nod factor treatment (Jardinaud et al., 2016).

IRON MAN peptides have been shown to be involved in iron transport in Arabidopsis (Grillet et al., 2018). In non-nodulating root samples IRON MAN genes were found to be expressed at nearly undetectable levels, but in nodulating roots all fourteen genes were actively expressed in the AON mutants by 48 to 72 hours after inoculation with nine of them up in wild type as well. Based on the timing of their induction, it would follow that these genes may be required for signaling root tissue to rapidly synthesize leghemoglobin, which keeps the oxygen tension in the nodule low enough for the bacterial nitrogenase to function.

### 4.2 Rhizobial response in AON mutants includes induction of TML genes

Our analysis of expression of functionally validated symbiotic nitrogen fixation genes (Roy et al., 2020) showed that the majority (over 70%) did not change expression in root tissues during the establishment of nodules. This is not surprising given that many of the genes encode receptors, enzymes, or transporters that may be constitutively expressed or encode proteins that are increased later in nodule development. However, 40 of the 206 genes did have a transcriptional response to rhizobia in wild type, and all these were also induced in both *sunn-4* and *rdn1-2*. An additional 16 genes had detectable increases following inoculation only in the AON mutants, perhaps because of the higher number of nodules forming on those roots.

Among the genes induced by rhizobia in both wild type and the AON mutants were *TML2* and *TML1*, encoding two related Kelch-repeat F-box proteins involved in suppressing nodulation (Gautrat et al., 2019). Both genes are targets of miR2111 (Gautrat et al., 2020). It has been proposed that miR2111 transported from shoots to roots maintains nodulation competence by keeping levels of *TML* mRNAs in roots low and that, following nitrate induction of *MtCLE35* (Moreau et al., 2021) or rhizobial induction of *MtCLE13* (Gautrat et al., 2020), a SUNN-dependent decrease in miR2111 expression allows accumulation of *TML* transcripts, proposed to halt further nodule development. Although increased *TML2* and *TML1* gene expression in roots following inoculation with rhizobia is expected in wild type, the finding that both genes were induced in a similar manner in the AON mutants early in nodulation is unexpected. The observed wild type patterns of transcript accumulation for both these genes in the *sunn-4* mutant demonstrates that early (24-48 hpi) accumulation is not SUNN-dependent. The SUNN-dependent increase in *TML2* and *TML1* transcript levels proposed from overexpression experiments with MtCLE35 in transgenic hairy roots at 14dpi (Moreau et al., 2021) did not measure transcript abundance of the *TML* genes, and measurement of *TML1* and *TML2* abundance in inoculated *sunn* mutants in (Gautrat et al., 2020) were performed in one branch of a split root at 5dpi. Our data gathered from a much earlier response to rhizobia in a single root adds new information to incorporate into the model with further experiments. A recent review noted that other factors in addition to SUNN may contribute to changes in *TML* levels (Okuma and Kawaguchi, 2021) and our data support this observation.

## 5. Conclusion

Transcriptome analysis demonstrates that expression of only a small set of genes is constitutively altered in the hypernodulating mutant *sunn-4*, despite the short root phenotype and auxin transport differences reported for *sunn* mutants. Of genes induced in nodulating roots of wild type, only a single gene, from a flavonoid synthesis pathway, was found to have a weaker response in *sunn-4*. The early rhizobial response of *sunn-4* included the wild type induction of both *TML2* and *TML1*, suggesting that early increased expression of these nodulation regulation genes is not sufficient for autoregulation.

## Supporting information

Supplemental Data Set 1

Supplemental Data Set2

## Author contributions

Elise Schnabel: Conceptualization, Methodology, Investigation, Validation, Writing - Original Draft. Suchitra Chavan: Investigation. William Poehlman: Methodology, Software. Yueyao Gao: Formal analysis, Visualization. F. Alex Feltus: Methodology, Writing - Review & Editing. Julia Frugoli: Conceptualization, Writing - Review & Editing.

## Declaration of competing interest

The authors have declared that no competing interests exist for this research work.

## Acknowledgements

This work is supported by NSF IOS 1444461 to Frugoli and Feltus and NSF IOS 2139351(Frugoli co-PI). The work was made possible in part, with support from the Clemson University Genomics and Bioinformatics Facility, which receives support from an Institutional Development Award (IDeA) from the National Institute of General Medical Sciences of the National Institutes of Health under grant number P20GM109094. The funders had no involvement in study design; in the collection, analysis and interpretation of data; in the writing of the report; and in the decision to submit the article for publication, beyond approval of the proposal for funding. Much of the computation was performed on the Clemson University Palmetto Cluster.

## Notes

### Competing Interest Statement

The authors have declared no competing interest.

