## Supplemental Fig 1 for "Transcriptome analysis of *Medicago truncatula* Autoregulation of Nodulation mutants reveals that disruption of the SUNN pathway causes constitutive expression changes in a small group of genes, but the overall response to rhizobia resembles wild type, including induction of *TML1* and *TML2*"

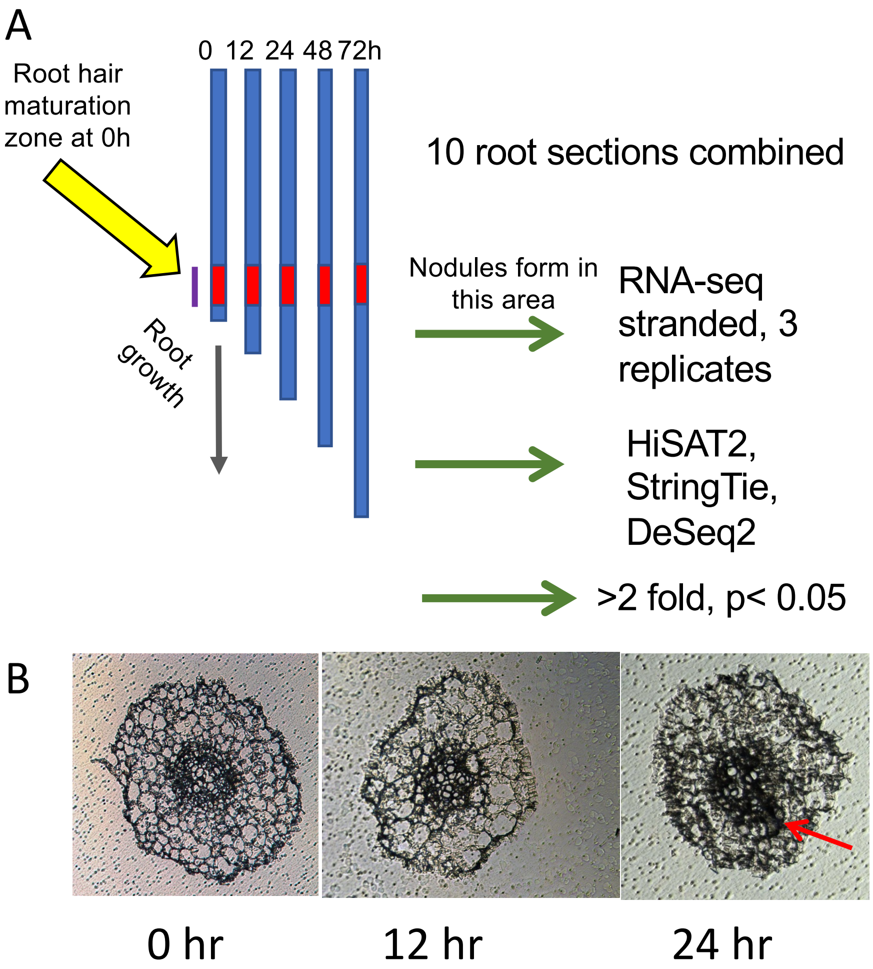


**Supplemental Figure 1. (A) Diagram of experimental procedure designed to increase signal to noise ratio** Twenty plants per genotype (t = 0 h) were collected and rhizobia added to the growth apparatus immediately after collection of the 0 h samples. Additional samples of 20 plants per genotype were collected at 12, 24, 48, and 72 h after the 0 h samples. Ten plants from each collection were used to determine average root length and 2 cm segments representing the zone of development of the first nodules were collected from the remaining 10 plants. At 0 h this region started 1 cm from the root tip, where the first full length root hairs were present. At later time points, this region was determined by calculating the average root growth since $t$=0 and adding this distance to 1 cm. (B) cross sections of roots harvested during first 3 time points. Cell division for nodule formation was occasionally observed at 24 hours (red arrow).
