## Supplemental Fig 2 for "Transcriptome analysis of *Medicago truncatula* Autoregulation of Nodulation mutants reveals that disruption of the SUNN pathway causes constitutive expression changes in a small group of genes, but the overall response to rhizobia resembles wild type, including induction of *TML1* and *TML2*"

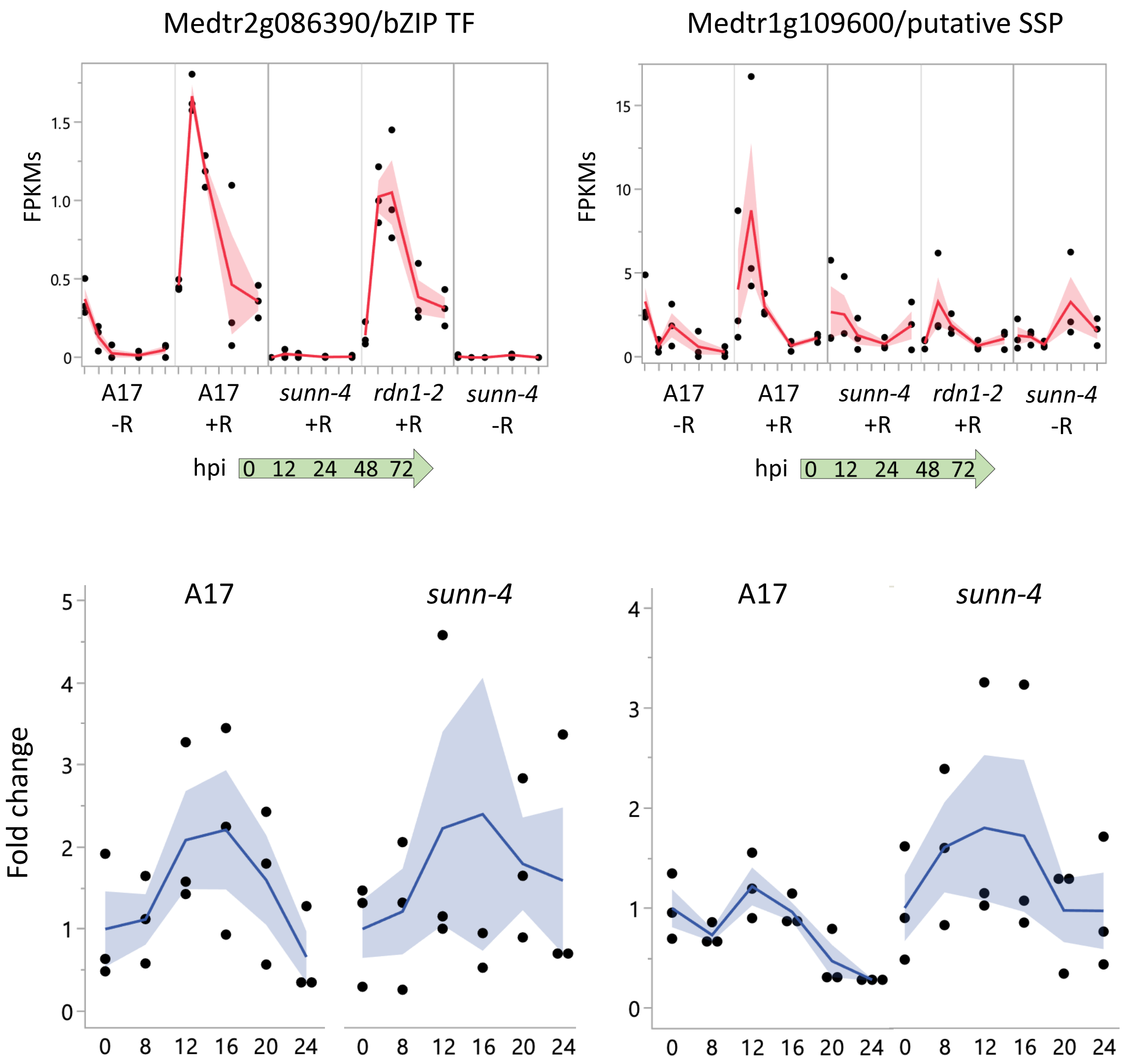


Supplemental Fig. 2. Induction of Medtr2g086390 and Medtr1g109600 in nodulating wild type roots was found by 12 hours post inoculation (hpi) in RNAseq data but was not verified as statistically significant by qPCR analysis (Kruskal-Wallis test with Bonferroni correction). FPKMs (black dots) and means (red lines) from three biological replicates are shown in the upper panels for time points 0 through 72 hpi for uninoculated (-R; wild type A17 and *sunn-4*) and inoculated (+R; wild type, *sunn-4*, and *rdn1-2*) root segments. Lower panels show expression over the first 24 hours post inoculation detected by qPCR (biological replicates as black dots; means as blue lines). Shading shows the standard error of the mean.
