## Supplemental Fig 3 for "Transcriptome analysis of *Medicago truncatula* Autoregulation of Nodulation mutants reveals that disruption of the SUNN pathway causes constitutive expression changes in a small group of genes, but the overall response to rhizobia resembles wild type, including induction of *TML1* and *TML2*"

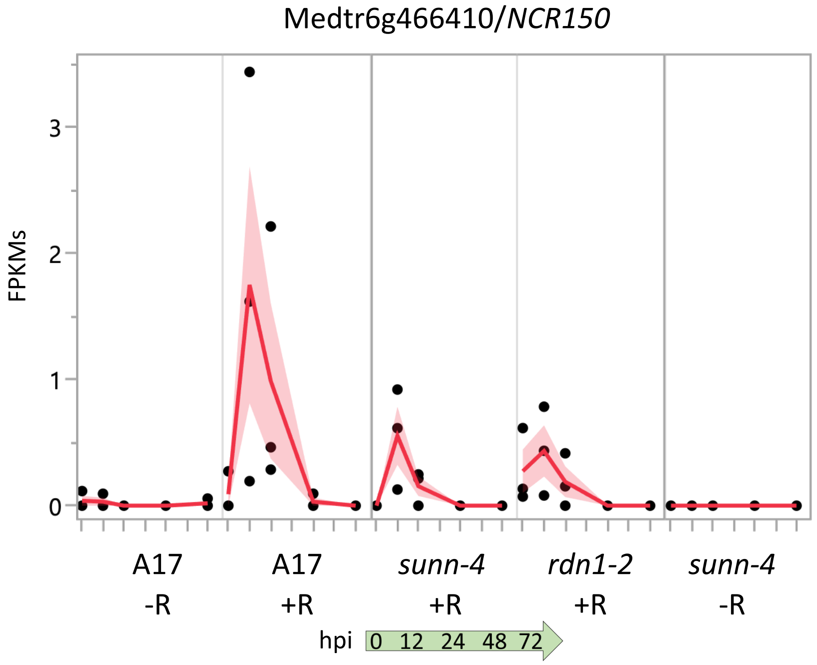


Supplemental Fig. 3. Transient induction of NCR150 (Medtr6g466410) at early time points following rhizobial inoculation. The graph shows FPKMs (black dots) and their means (red lines) over time from RNAseq. Shading is the standard error of the mean.
