## Supplemental Fig 4 for "Transcriptome analysis of *Medicago truncatula* Autoregulation of Nodulation mutants reveals that disruption of the SUNN pathway causes constitutive expression changes in a small group of genes, but the overall response to rhizobia resembles wild type, including induction of *TML1* and *TML2*"

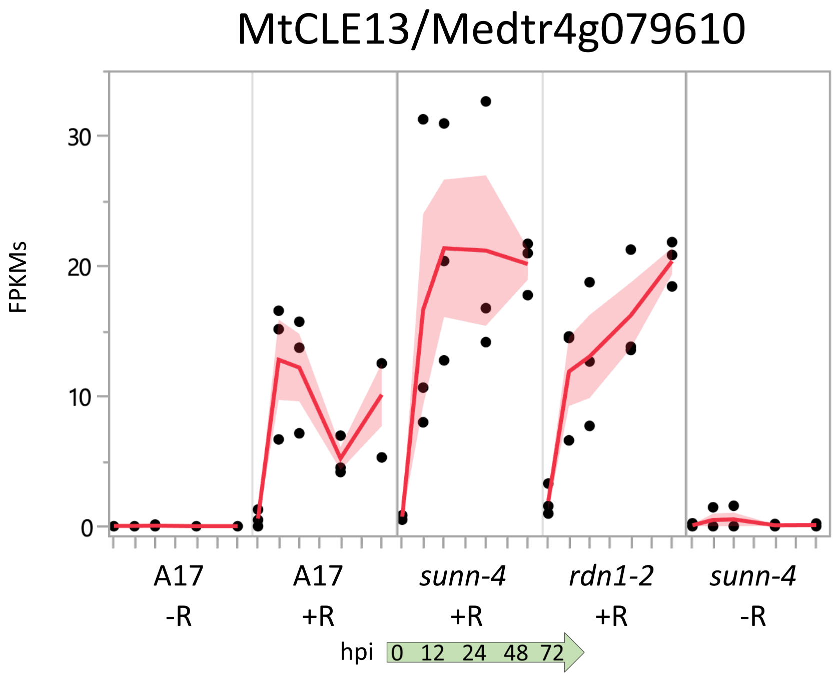


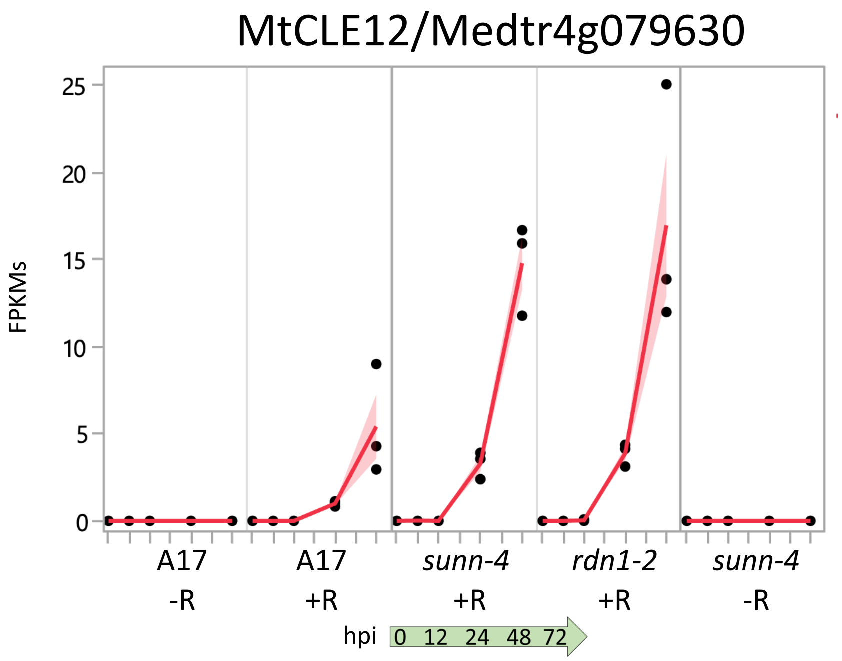


Supplemental Fig. 4. Induction of *MtCLE13* and *MtCLE12* following rhizobial inoculation. The graph shows FPKMs (black dots) and their means (red lines) from RNAseq. Shading is the standard error of the mean.
